# Comprehensive analysis of miRNA and protein profiles within exosomes derived from canine lymphoid tumour cell lines

**DOI:** 10.1101/476309

**Authors:** Hajime Asada, Hirotaka Tomiyasu, Takao Uchikai, Genki Ishihara, Yuko Goto-Koshino, Koichi Ohno, Hajime Tsujimoto

## Abstract

Exosomes are small extracellular vesicles released from almost all cell types, which play roles in cell-cell communication. Recent studies have suggested that microenvironmental crosstalk mediated by exosomes is an important factor in the escape of tumour cells from the anti-tumour immune system in human haematopoietic malignancies. Here, we conducted comprehensive analysis of the miRNA and protein profiles within the exosomes released from four canine lymphoid tumour cell lines as a model of human lymphoid tumours. The results showed that the miRNAs and proteins abundantly contained in exosomes were similar among the four cell lines. However, the profiles of miRNA within exosomes differed among the cell lines and reflected the expression pattern of miRNAs of the parent cells. In the comparison of the amounts of miRNAs and proteins among the cell lines, those of three miRNAs (miR-151, miR-8908a-3p, and miR-486) and CD82 protein differed between exosomes derived from vincristine-sensitive and resistant cell lines. Further investigations are needed to elucidate the biological functions of the exosomal contents in the microenvironmental crosstalk of lymphoid tumours.

## Introduction

Exosomes are small extracellular vesicles released from almost all cell types, including immune cells and tumour cells [1], as the intracellular endosome component. Although exosomes were initially considered cellular waste, they have been shown to contain various molecules from the original cells, including proteins, functional mRNAs and miRNAs, and deliver these biological messages into the recipient cells [1,2]. To date, it has also been reported that tumour cells release a number of exosomes and they stimulate tumour cell growth and modify the immune cell response to promote tumour progression and metastasis in several human tumors, including colorectal cancer [3], breast cancer [4], melanoma [5], and pancreatic cancer [6]. Thus, the interaction between tumour cell-derived exosomes and recipient cells in the microenvironment of solid tumours is considered an important factor in tumour progression, metastasis, cell survival, and escape from the anti-tumour immune system.

Exosomes have also been suggested to play important roles in the microenvironmental crosstalk of human haematopoietic tumours, including leukaemia and lymphoma [7,8]. It has been reported that exosomes derived from acute/chronic myeloid leukaemia and lymphoma cells inactivate natural killer cells and suppress the anti-tumour immune response [7–9]. In addition, exosomes have been reported to be associated with drug resistance in these tumours [7]. For instance, it was reported that exosomes derived from imatinib-resistant chronic leukaemia cells could confer imatinib-resistance traits into sensitive cells by delivering miR-365 [10]. It was also reported that exosomes derived from bone marrow stromal cells decreased the sensitivity of acute lymphoblastic leukaemia cells to etoposide [11]. Based on this background information, it has been considered that studies on the molecules contained in exosomes released from haematopoietic tumour cells could provide insight into the pathophysiology of these tumours. Although the miRNA profile within exosomes was reported in Gamma-Herpesvirus-infected lymphoma cell lines [12] and lymphocytic leukaemia cells [13], no study has yet comprehensively analysed the miRNA and protein profiles of exosomes derived from haematopoietic malignancies.

Lymphoma is a haematopoietic malignancy originating from lymphoid cells, and it is categorised into more than 80 distinct subtypes [14]. Among them, Non-Hodgkin lymphoma (NHL) is the most common type of lymphoma in humans and dogs [15]. It has been reported that canine lymphoma shares many characteristics of human NHL, including clinical presentation, immunophenotypic composition, chemotherapeutic protocols, and response to treatment [15,16]. Therefore, canine lymphoma has been advocated as an ideal model for studying human NHL [15,16].

The aim of this study was to comprehensively analyse the miRNA and protein profiles within the exosomes released from canine lymphoid tumour cells.

## Results

### Exosome isolation and preparation of total RNA of exosomes and parent cells

The size distributions of exosomes isolated from four canine lymphoid tumour cell lines, CLBL-1, GL-1, UL-1, and Ema, are shown in S1 Fig. The average size was between approximately 100–150nm in each cell line. The RNA integrity numbers (RINs) and size distributions of total RNA samples taken from exosomes and parent cells are shown in S2 Fig. Although there were common peaks corresponding to ribosomal RNAs in exosomal RNA of the four cell lines, the distributions of RNA sizes were clearly different between exosomes and parent cells.

### Exosomal miRNA profiles

At first, the miRNA profiles of exosomes and parent cells were investigated via small RNA sequencing analysis. A minimum of 20 million raw reads were generated for each sample (see S1 Table). The number of reads mapped to miRNA and the mapping rate to miRNA was comparatively lower in Ema than the other three cell lines. Therefore, data for Ema were omitted in the statistical comparison of the quantities of miRNAs among cell lines using small RNA sequencing data.

Then, hierarchical clustering using the amounts of miRNA in CLBL-1, GL-1, and UL-1 was conducted. This analysis yielded three clusters composed of exosomes and parent cells of each cell line (Fig 1a). In addition, in the PCA plots, exosomes and cells clustered similarly for each cell line (Fig 1b). The results of these analyses were similar when the data from Ema were included (see S3 Fig).

**Fig 1. Hierarchical clustering (a) and PCA plots (b) for miRNA profiles of exosomes and parent cells of CLBL-1, GL-1, and UL-1.** Exosomes and parent cells clustered similarly for each cell line and the profiles were different among cell lines. Orange dots (exosomes) and red dots (parent cells) correspond to CLBL-1, violet dots (exosomes) and blue dots (parent cells) to GL-1, and grey dots (exosomes) and black dots (parent cells) to UL-1.

The top ten miRNAs contained in exosomes and parent cells are listed in Table 1. Among these miRNAs, five miRNAs (let-7f, let-7g, miR-7, miR-30d, and miR-92a) were commonly contained in exosomes and cells of the four cell lines.

**Table 1.** The top 10 miRNAs abundantly contained in the exosomes and parent cells in this study.

| CLBL-1 |  | GL-1 |  | UL-1 |  | Ema |  |
| --- | --- | --- | --- | --- | --- | --- | --- |
| Exosome | Parent cell | Exosome | Parent cell | Exosome | Parent cell | Exosome | Parent cell |
| miR-148a | miR-148a | miR-148a | miR-148a | miR-7 | miR-7 | miR-7 | let-7g |
| miR-7 | let-7g | let-7f | let-7g | miR-378 | miR-99a | let-7g | miR-7 |
| let-7g | miR-363 | let-7g | miR-10a | miR-99a | miR-378 | let-7f | miR-363 |
| let-7f | miR-7 | miR-30d | let-7f | miR-30d | miR-30d | miR-30d | let-7f |
| miR-146a | miR-99a | miR-10a | miR-30d | let-7g | let-7g | miR-363 | miR-30d |
| miR-30d | miR-30d | miR-378 | miR-378 | let-7f | miR-10a | miR-21 | miR-21 |
| miR-99a | miR-128 | miR-7 | miR-7 | miR-363 | miR-363 | miR-148a | miR-128 |
| miR-20a | miR-92a | let-7a | miR-128 | miR-10a | miR-128 | miR-26a | miR-26a |
| miR-378 | let-7f | miR-103 | miR-21 | miR-92a | let-7f | miR-155 | miR-92a |
| miR-92a | miR-146a | miR-92a | let-7a | miR-103 | miR-25 | miR-92a | miR-155 |

In the comparison of the amounts of miRNAs between cells and exosomes, the amounts of 39, 20, and 24 miRNAs were significantly different in CLBL-1, UL-1, and Ema, respectively (q < 0.01) (Fig 2). Among these miRNAs, the amount of miR-350 was significantly higher in exosomes than parent cells in all the three cell lines, and those of miR-22, miR-671, and miR-8865 were significantly lower in exosomes than parent cells in these cell lines (see S4 Fig). On the other hand, no miRNA displayed a significant difference in amount between exosomes and cells in GL-1.

**Fig 2. Heat maps showing the miRNAs whose amounts were significantly different between exosomes and parent cells of CLBL-1 (a), UL-1 (b), and Ema (c).** The amounts of 39, 20, and 24 miRNAs were significantly different in CLBL-1, UL-1, and Ema, respectively (q < 0.01).

The difference in the amount of miR-350 between exosomes and parent cells was confirmed by RT-qPCR (Fig 3). However, the amounts of miR-22, miR-671, and miR-8865 were not significantly different between exosomes and parent cells according to RT-qPCR. Following quantitative analysis, prediction of target genes was conducted for miR-350 using miRbase, and the top 10 target genes of the miRNA were extracted (see S2 Table). These target genes of miR-350 did not include those previously reported to be associated with the pathophysiology of tumour cells.

**Fig 3. Comparison of the amounts of miR-350 (a), miR-671 (b), miR-22 (c), and miR-8865 (d) between exosomes and parent cells in the four cell lines.** The amount of miR-350 is significantly different between exosomes and parent cells, whereas those of miR-22, miR-671, and miR-8865 were not significantly different. All data represent the mean ± SD of three independent experiments. *P < 0.05.

### Exosomal protein profiles

The results of separating exosomal proteins from each cell line by SDS-PAGE are shown in S5 Fig. Exosomal protein profiles were investigated by liquid chromatography-tandem mass spectrometry (LC-MS/MS). This analysis identified a total of 1,890 proteins among peptides extracted from exosomes of the four cell lines.

The top twenty proteins that were detected in each cell line by LC-MS/MS are listed in Table 2. As is the case with miRNAs, 13 proteins were commonly contained in the four cell lines. The abundantly contained proteins included those related to the cytoskeleton (β-actin and tubulins) and heat shock proteins. Except for these abundant proteins, CD63 was detected in exosomes of all four cell lines and CD81 was detected in those of CLBL-1, GL-1 and UL-1 among the exosome marker proteins, although CD9 was not detected in any cell lines.

**Table 2.** The top 20 proteins abundantly contained in exosomes in this study.

| CLBL-1 | GL-1 | UL-1 | Ema |
| --- | --- | --- | --- |
| ACTB | ACTB | ACTB | ACTB |
| TUBB | TUBB | TUBB | TUBB |
| TUBB4B | TUBB4B | TUBB4B | TUBB4B |
| TUBB2B | TUBB2B | TUBB2B | TUBB2B |
| TUBB4A | TUBB4A | TUBB4A | TUBB4A |
| TUBA1C | TUBA1C | TUBA1C | TUBA1C |
| TUBA4A isoform X1 | TUBA4A isoform X1 | TUBA4A isoform X1 | TUBA4A isoform X1 |
| FLNA | FLNA | FLNA | FLNA |
| TLN1 isoform X4 | TLN1 isoform X4 | TLN1 isoform X4 | TLN1 isoform X4 |
| MYH9 | MYH9 | MYH9 | MYH9 |
| FAS | FAS | FAS | FAS |
| EEF2 | EEF2 | EEF2 | EEF2 |
| ACLY isoform X1 | ACLY isoform X1 | ACLY isoform X1 | ACLY isoform X1 |
| HSP90B | NCL | HSP90B | HSP90B |
| CCT2 isoform X1 | IQGAP1 | CCT2 isoform X1 | CCT2 isoform X1 |
| TUBBA3 | DYNC1H1 | DYNC1H1 | DYNC1H1 |
| CCT8 isoform X2 | CLTC isoform X1 | CLTC isoform X1 | CLTC isoform X1 |
| CCT8 isoform X1 | GAPDH | GAPDH | GAPDH |
| CENP | CENP | EEF1A1 | EEF1A1 |
| HSP71 | HSP71 | EPRS isoform X1 | PKM isoform X1 |

### Comparison of exosomal miRNA and protein profiles between vincristine sensitive (VCR-S) cell lines and vincristine resistant (VCR-R) cell lines

The exosomal miRNA profiles were also compared between the VCR-S cell lines (CLBL-1 and GL-1) and the VCR-R cell line (UL-1) (see S6 Fig). In data from small RNA sequencing, the amounts of 11 miRNAs within exosomes were significantly lower in VCR-S cell lines than in the VCR-R cell line, and those of 5 miRNAs were significantly higher in VCR-S cell lines than in the VCR-R cell line (q < 0.01). In parent cells, the amounts of 8 miRNAs were significantly lower in VCR-S cell lines than in the VCR-R cell line, and those of 7 miRNAs were higher in VCR-S cell lines than in the VCR-R cell line (q < 0.01).

Among these miRNAs, the significant differences in the amounts of miR-151, miR-8908a-3p, and miR-486 were confirmed by RT-qPCR using the four cell lines including Ema, which is resistant to VCR (Fig 4). The amounts of miR-151 and miR-8908a-3p within exosomes and parent cells in VCR-S cell lines were significantly lower than those in VCR-R cell lines (P < 0.01). The amount of miR-486 within exosomes and parent cells in VCR-S cell lines was significantly higher than those in VCR-R cell lines (P < 0.01). The target genes were predicted for miR-151, miR-8908a-3p, and miR-486, and the top 10 target genes of each miRNA were extracted (see S2 Table). These genes included those that have been reported to be associated with the biological behaviour of tumour cells (*NTRK2*, *MAPK8*, *BCOR*, and *PIK3R1* genes).

**Fig 4. Comparison of the amounts of miR-151 (a), miR-8908a-3p (b), and miR-486 (c) between VCR-S and VCR-R cell lines.** The amounts of miR-151 and miR-8908a-3p in VCR-S cell lines were significantly lower than in VCR-R cell lines, and that of miR-486 in VCR-S cell lines were significantly higher than in VCR-R cell lines. All data represent the mean ± SD of three independent experiments. *P < 0.05.

Following the LC-MS/MS analysis, proteins that were detected only in VCR-S or VCR-R cell lines were also extracted (Table 3). Among these proteins, the difference in the amount of CD82 was validated by western blotting (Fig 5). CD82 was detected in exosomes of CLBL-1 and GL-1, while no band corresponding to CD82 was detected in exosomes of UL-1 and Ema. This protein was not detected in parent cells of all the four cell lines. HSP90B, which was selected as a protein that is abundantly contained in exosomes, was detected in exosomes from all four of the cell lines.

**Table 3.**
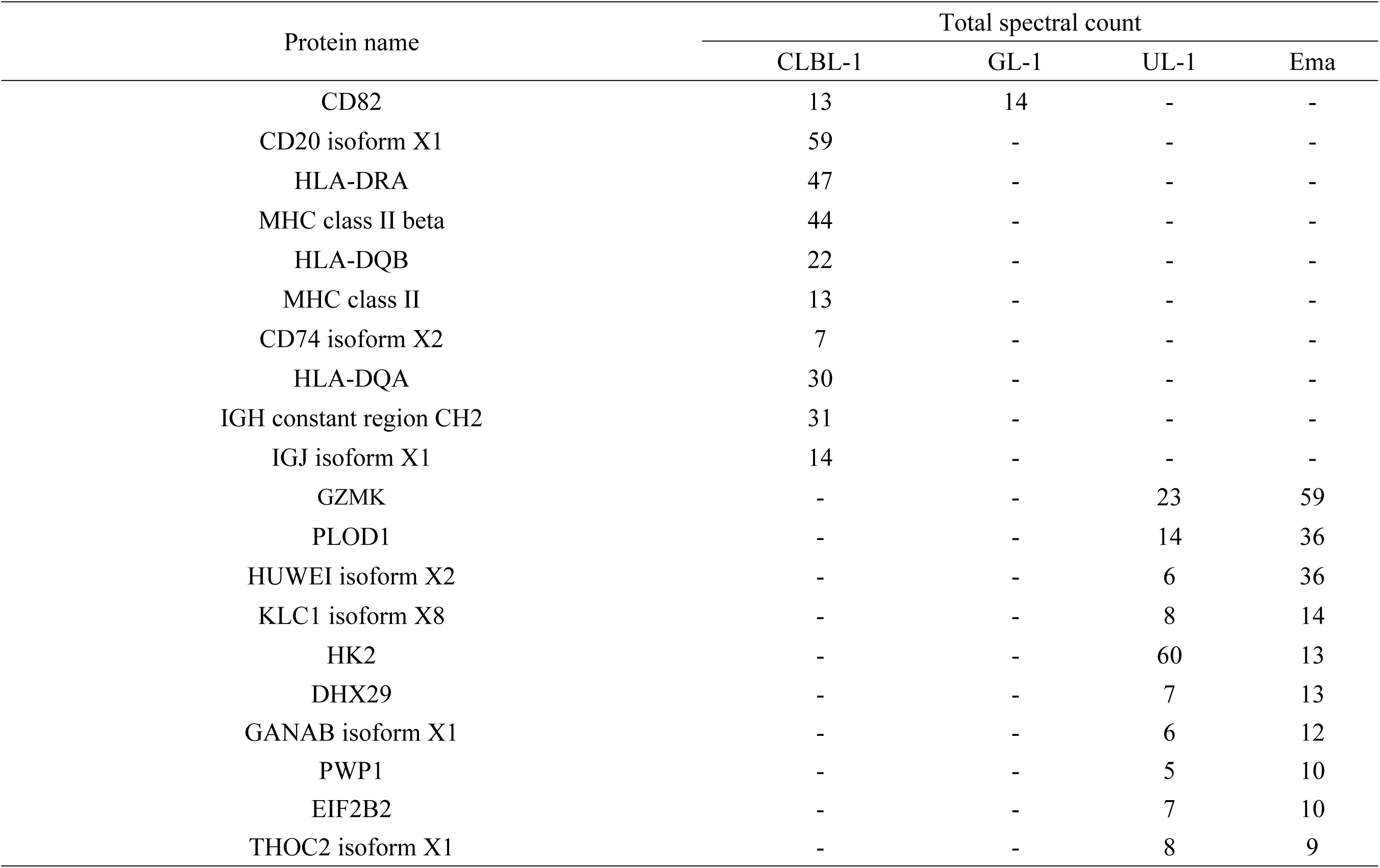
Exosomal proteins detected only in vincristine-sensitive or vincristine-resistant cell lines.

**Fig 5. Western blotting for CD82 using proteins extracted from exosomes (a) and parent cells (b) of each cell line.** HSP90B and β-actin were selected for internal control for exosomes and parent cells, respectively. CD82 protein is detected in the exosomes of CLBL-1 and GL-1, whereas it was not detected in parent cells in any of the four cell lines. The figures of detection of CD82 within exosomes and parent cells were cropped from the different parts of the same figure of the membrane. The figures of HSP90B and β-actin were cropped from the figures of the different membrane. The full-length figures of blotting membrane are shown in S7 Fig.

## Discussion

In the present study, the miRNA and protein profiles within exosomes derived from four canine lymphoid tumour cell lines were comprehensively analysed by small RNA sequencing and LC-MS/MS.

In small RNA sequencing, the mapping rate of the reads to canine miRNA was comparatively lower in both exosomes and parent cells of Ema than the other three cell lines. In the hierarchical clustering analysis and PCA plots for three cell lines, three distinct clusters composed of the exosomes and parent cells of each cell line were observed. Therefore, it was indicated that miRNA profiles within exosomes reflect those of parent cells and the profiles of exosomal miRNA varied among cell lines.

Small RNA sequencing also revealed that five miRNAs (let-7f, let-7g, miR-7, miR-30d, and miR-92a) were abundantly contained both in exosomes and parent cells of all four of the cell lines. Previous studies have reported that exosomes derived from tumour cells contain miRNAs of the let-7 family [17,18]. It has also been reported that miR-30d and miR-92a are abundant in exosomes of Gamma-Herpesvirus-infected lymphoma cell lines [12]. Therefore, these miRNAs might be associated with the pathophysiology of lymphoid tumours, and further studies are needed to reveal the biological roles of these miRNAs in exosomes derived from tumour cells.

In the comparison of the amounts of miRNAs between exosomes and parent cells, significant differences were observed for 39, 20, and 24 miRNAs in CLBL-1, UL-1, and Ema, respectively, whereas there was no significant difference in GL-1. Among these miRNAs, the significant differences in the amounts of miR-350 between exosomes and parent cells were confirmed in all four cell lines by RT-qPCR. The predicted target genes of miR-350 did not include those previously reported to be associated with the pathophysiology of tumour cells. However, miR-350 was reported to promote apoptosis through down-regulation of *PIK3R3* gene [19]. Further studies are needed to reveal the functions of miR-350 in the microenvironmental crosstalk in lymphoma.

LC-MS/MS analysis revealed that exosomes derived from each cell line contain various types of protein. Most of the proteins abundantly contained in exosomes were common among the four cell lines, including those related to cytoskeleton, such as β-actin, tubulins, and heat shock proteins. In addition, CD63 or CD81 were also detected in exosomes derived from each cell line. Exosomal markers have been reported to include members of the tetraspanin family (CD9, CD63, and CD81) and heat shock proteins (HSP60, HSP70, and HSP90) [20,21]. It was also reported that exosomes derived from Jurkat cells contain β-actin and tubulins [22]. Thus, those results in previous studies are consistent with those in the present study.

In the comparison of the amounts of miRNA within the exosomes, the amounts of miR-151, miR-8908a-3p, and miR-486 were confirmed to be different between VCR-S cell lines and VCR-R cell lines by RT-qPCR. The amounts of miR-151 and miR-8908a-3p were significantly lower in VCR-S cell lines, while miR-486 was significantly more abundant in these cell lines. The target genes of miR-151 included the gene *NTRK2*, a member of neurotrophic tyrosine receptor kinase family. The expression of *NTRK2* was reported to be down-regulated in patients with breast cancer with a poor prognosis [23]. The expression of this gene was also reported to suppress anoikis by activating the PI3K/Akt pathway in human ovarian cancer cells [24]. The target genes of miR-8908a-3p included *MAPK8* (also known as *JNK1*) and *BCOR*. The *MAPK8* gene is a member of the MAP kinase and JNK family, and involved in various cellular processes including cell proliferation, differentiation, and apoptosis [25,26]. The *BCOR* gene encodes a co-repressor of BCL6, a transcriptional repressor that is required for formation of germinal centres [27,28] and silences various genes involved in the cell cycle and apoptosis [29]. The target genes of miR-486 included *PIK3R1*, one of the oncogenes that promotes cell proliferation and tumour cell invasion [30]. Based on this evidence, it is possible that these miRNAs might be associated with the resistance to VCR in lymphoid tumours. Further studies are needed to elucidate the association of these miRNAs with drug resistance and microenvironmental crosstalk in lymphoid tumours.

Among the proteins detected by LC-MS/MS in the present study, CD82 were detected in the exosomes of VCR-S cell lines but not in those of VCR-R cell lines, and the difference in its amount was confirmed by western-blotting. In addition, CD82 was not detected in proteins extracted from parent cells of CLBL-1 and GL-1, suggesting that CD82 was selectively delivered into exosomes in these cell lines. CD82 has been reported to suppress tumour metastasis [31] and be associated with tumour cell growth [32] and survival [33]. Therefore, it is possible that CD82 expression in exosomes might be associated with the biological behaviour of tumour cells, including metastasis, tumour growth, cell survival, and drug sensitivity, via its function in microenvironmental crosstalk in lymphoid tumours. Further studies are needed to investigate the biological roles of CD82 in microenvironmental crosstalk in lymphoid tumours.

In conclusion, most of the miRNAs and proteins abundantly contained in exosomes are common among the four cell lines, but the miRNA profiles in exosomes reflect those of parent cells and differ among cell lines. In addition, miR-151, miR-8908a-3p, miR-486, and CD82 proteins were differentially abundant within the exosomes between VCR-S and VCR-R cell lines. Further investigations are needed to elucidate the biological functions of these molecules in the crosstalk between tumour cells and tumour microenvironment.

## Materials and methods

### Cell lines and cell culture

Four canine lymphoid tumour cell lines (CLBL-1, GL-1, UL-1, and Ema) were used in this study: CLBL-1, a canine B-cell lymphoma cell line [34]; GL-1, a canine B-cell leukaemia cell line [35]; UL-1, a canine T-cell lymphoma cell line [36]; and Ema; a canine T-cell lymphoma cell line [37]. UL-1 and Ema were established from dogs with lymphoma showing drug resistance after chemotherapy, whereas CLBL-1 and GL-1 were established from dogs with leukaemia or lymphoma who were not subjected to chemotherapy. CLBL-1, GL-1, and Ema were kindly provided by Dr. Rütgen, University of Veterinary Medicine Vienna, Austria, Dr. Nakaichi, Yamaguchi University, Japan, and Dr. Mizuno, Yamaguchi University, Japan, respectively. Our group established UL-1 previously [36]. Our previous study reported that CLBL-1 and GL-1 were sensitive to vincristine, and UL-1 and Ema were resistant to vincristine [38]. These cell lines were cultured in RPMI-1640 medium at 37°C, with 10% foetal bovine serum (Biowest, Nuaille, France) in a humidified atmosphere containing 5% CO_2_.

## Exosome isolation and preparation of total RNA and protein of exosomes and parent cells

Exosomes were isolated from 3×10^7^ cells (CLBL-1, GL-1, and UL-1) and 2×10^7^ cells (Ema) cultured for 24h in growth medium without foetal bovine serum. Exosomes were isolated from cell culture media using the Total Exosome Isolation (from cell culture media) (ThermoFisher Scientific, Waltham, MA, USA), and exosome protein and RNA were prepared using the Total Exosome RNA and Protein Isolation Kit (ThermoFisher Scientific) according to the manufacturer’s instructions. The number and sizes of isolated exosomes were measured using NanoSight NS300 system (Malvern Instruments, Malvern, UK). The concentrations of exosome protein samples were measured using Micro BCA Protein Assay (ThermoFisher Scientific), and the concentrations and size distributions of exosomal RNA samples were measured using Agilent RNA 6000 Pico Kit and Agilent 2100 Bioanalyzer (Agilent Technologies, Palo Alto, CA, USA). Total RNA of each parent cell line was extracted using miRNeasy Mini Kit (QIAGEN, Limburg, Netherlands), and concentration and integrity were measured as described above. Each total RNA sample was prepared in duplicate.

### Small RNA sequencing and data processing

Small RNA sequencing libraries were prepared with 156 ng of total RNA using NEB Next Multiplex Small RNA Library Prep Kit (New England Biolabs, Ipswich, MA, USA). RNA sequencing was performed in duplicate using NextSeq500 (Illumina, San Diego, CA, USA) with High Output Kit (Illumina) as stranded, single 36-base reads following the manufacturer’s instruction.

Raw BCL data for each sample were de-multiplexed with bcl2fastq (version 2.18.0.12) and were stored in independent FASTQ files. The sequence data were trimmed with Trimommatic (version 0.36) [39] to clean up sequences with low-quality and those with sequencing adaptors. After trimming, a subset of short reads was aligned to cfa_MiR_453 (http://www.targetscan.org/) with Bowtie2 (version 2.2.9) [40]. The depth of the reads aligned to cfa_MiR_453 was quantified using Samtools (version 1.3.1). Counts per million (CPM) was imported into R (version 3.3.2) and principal component analysis was conducted. Then, miRNA counts for each sample were imported into R for differential expression analysis with EdgeR [41,42]. Cluster3.0 and Java Treeview (version 1.1.6r4) were used for hierarchical clustering and visualization. The data from small RNA sequencing in this study are available in the DDBJ Sequenced Read Archive database with the accession number DRA006696.

### Quantitative real-time RT-PCR

The amounts of miRNAs extracted from the small RNA sequencing data were validated by RT-qPCR using TaqMan MicroRNA Assay (Applied Biosystems, Foster City, CA, USA). The candidate miRNAs selected for validation are listed in the Supplementary Table S3 online. Briefly, 3.3 ng of total RNA was reverse transcribed using TaqMan MicroRNA Reverse Transcription Kit (Applied Biosystems), and qPCR was performed using TaqMan MicroRNA Assay and Thermal Cycler Dice Real Time System TP800 (Takara Bio, Shiga, Japan). Data were expressed as mean C_T_ values of three independent experiments performed in triplicate. C_T_ values were determined using the second derivative maximum method, in which a C_T_ value is expressed as the cycle number at which the second derivative was at its maximum. After validation, target genes of the miRNAs were predicted using miRbase (http://www.mirbase.org/) [43,44].

### LC-MS/MS

Protein profiles of exosomes were analysed by LC-MS/MS. An EASY-Spray column (15 cm × 75 µm I.D., 3 µm, ThermoFisher Scientific) was employed for separation of each exosomal protein sample at the flow rate of 300 nl/min. A quadrupole tandem mass spectrometer (Q Exactive Plus, ThermoFisher Scientific) was used in positive ion mode for analytic detection. The raw MS spectra data were queried against the NCBI Canine protein sequence database using the MASCOT database search engine, and peptides were quantified according to the spectral counts.

### Western-blotting

Expressions of the candidate proteins extracted from the LC-MS/MS data were verified by western blotting. One µg of protein extracted from exosome or parent cells was separated by SDS-PAGE and blotted onto a PVDF membrane. The membranes were blocked in 5% skimmed milk and incubated with primary antibodies against CD82, HSP90B, or β-actin. HSP90B was selected as the internal control of the exosomal protein. Then, the membranes were incubated with secondary antibodies. The antibodies, dilutions, and incubation temperatures are shown in the Supplementary Table S4 online. After incubation, positive immunoreactivity was detected using Luminata Forte Western HRP Substrate (Merck Millipore, Darmstadt, Germany) and visualized using a ChemiDoc XRS Plus (Bio-Rad Laboratories, Hercules, CA, USA).

### Statistical analysis

In the differential expression analysis using EdgeR, a false discovery rate (q-value) of less than 0.01 was considered statistically significant. One-way ANOVA followed by Tukey’s post-hoc test was performed for multiple comparisons of miRNA quantities in the RT-qPCR using the STATMATE (ATMS, Tokyo, Japan) software, and P-values of less than 0.05 were considered statistically significant.

## Supporting information

S1 Table. The mean numbers of raw reads, reads mapped to miRNA, and the mapping rates to miRNA.

S2 Table. The top 10 predicted target genes of miRNAs extracted in this study.

S3 Table. miRNAs extracted for validation by RT-qPCR and TaqMan MicroRNA Assay IDs for these miRNAs.

S4 Table. Primary and secondary antibodies for detection of CD82, HSP90B, and β-actin.

---

**S1 Fig.**
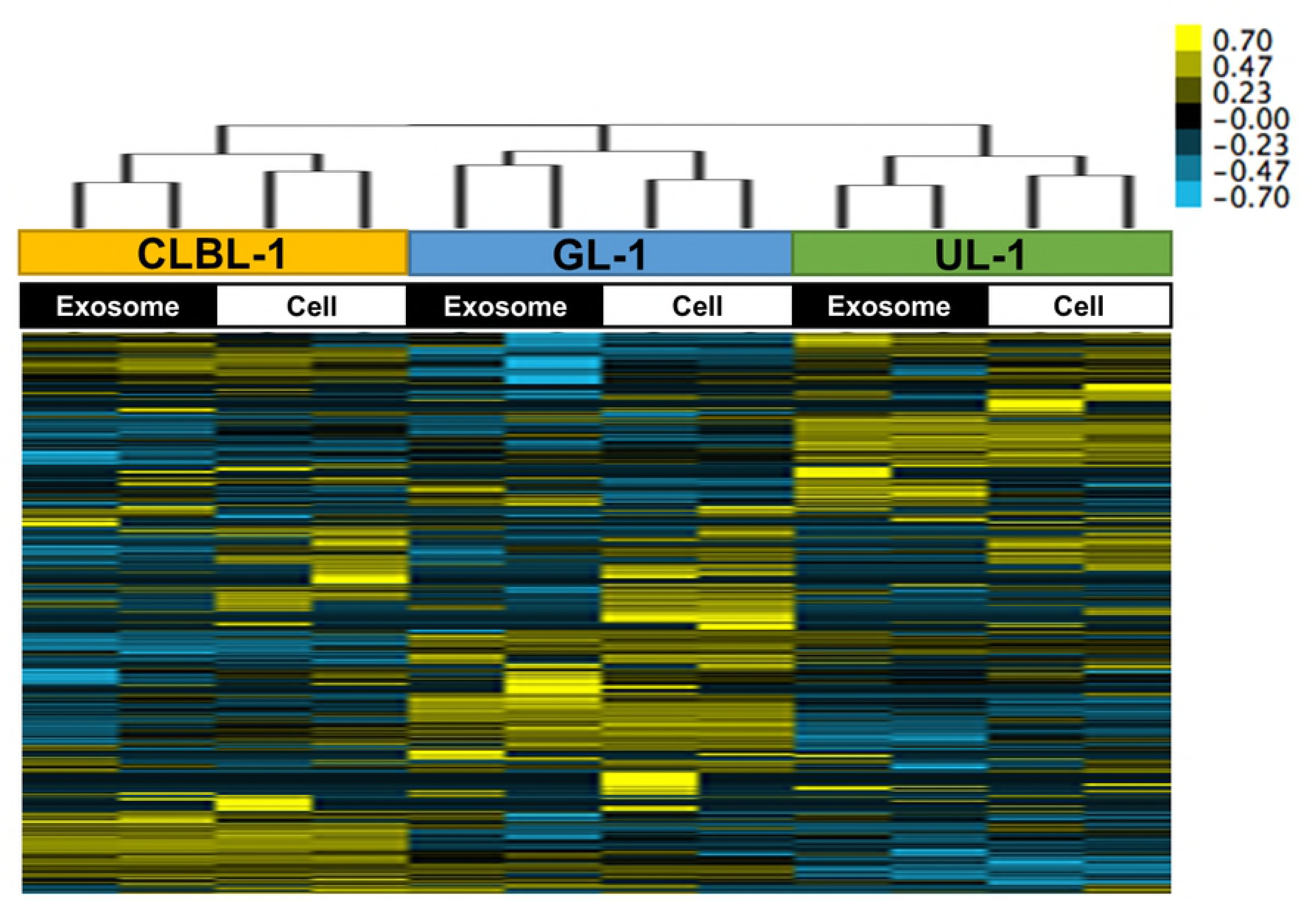
The size distributions of exosomes isolated from CLBL-1 (a), GL-1 (b), UL-1 (c), and Ema (d).

**S2 Fig.**
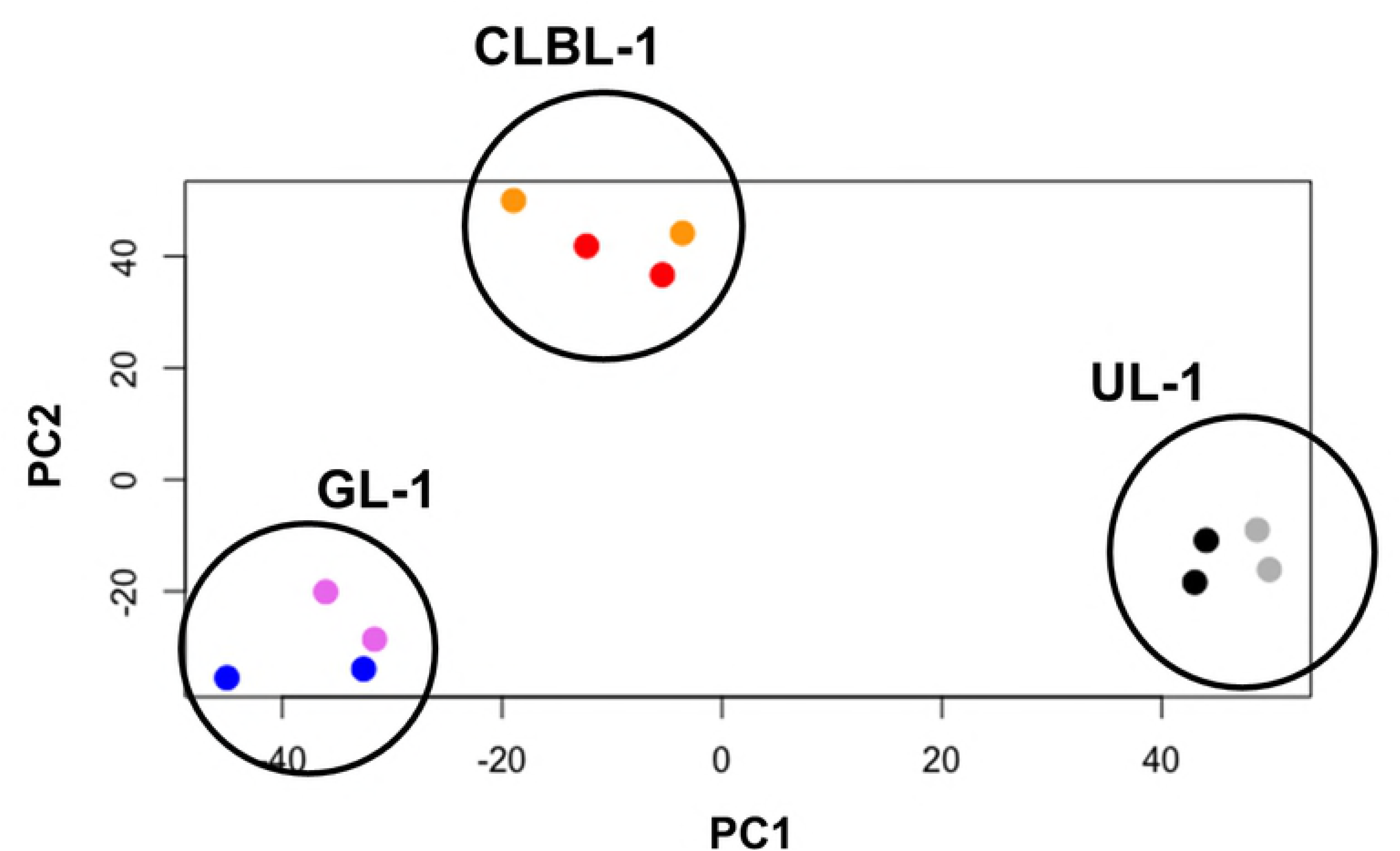
The RNA integrity numbers (RINs) and size distributions of total RNA samples derived from exosomes and parent cells in CLBL-1 (a), GL-1 (b), UL-1 (c), and Ema (d). “18S” indicates the peak of 18S ribosomal RNA, and “28S” indicates that of 28S ribosomal RNA.

**S3 Fig.**
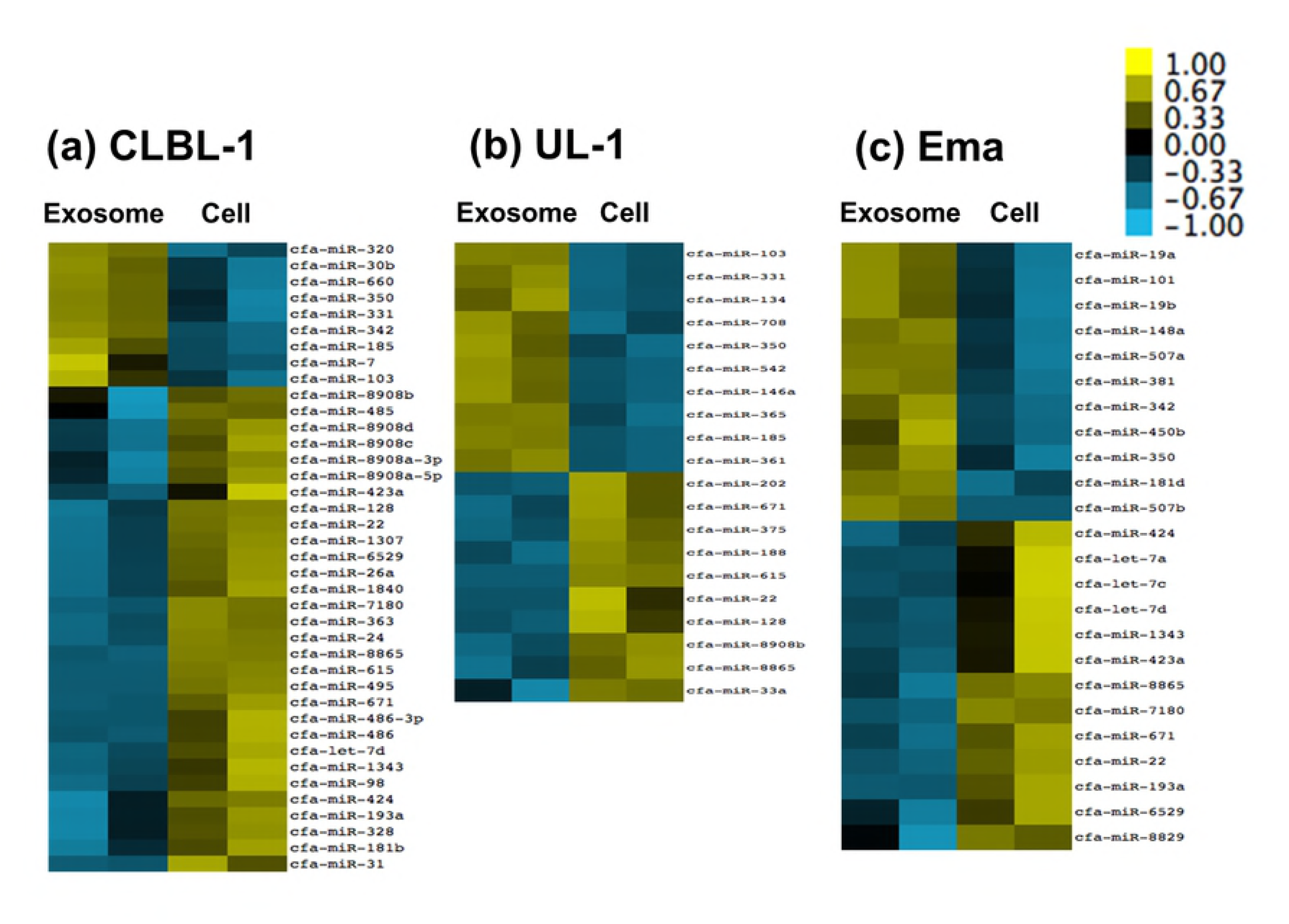
PCA plots analysis including data of Ema cell line. Exosomes and parent cells clustered similarly for each cell line and the profiles are different among cell lines. Orange dots (exosomes) and red dots (parent cells) correspond to CLBL-1, violet dots (exosomes) and blue dots (parent cells) to GL-1, grey dots (exosomes) and black dots (parent cells) to UL-1, and yellow dots (exosomes) and green dots (parent cells) to Ema.

**S4 Fig.**
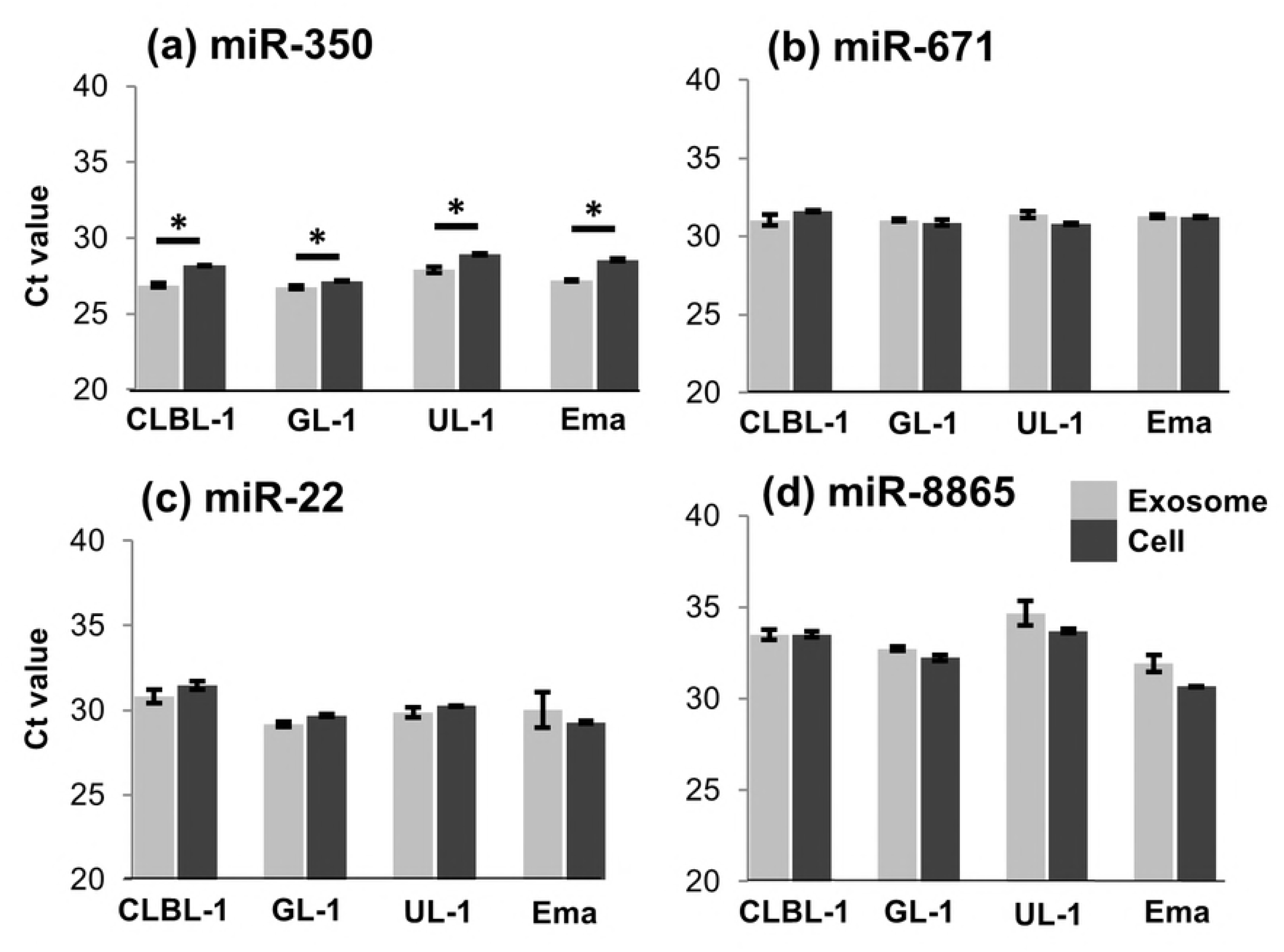
The Venn diagram showing the common miRNAs whose amounts were significantly different between exosomes and parent cells in small RNA sequencing. The names of miRNAs that were significantly more abundant in exosomes than parent cells are shown in red, and those that were significantly less abundant in exosomes than parent cells are shown in blue.

**S5 Fig.**
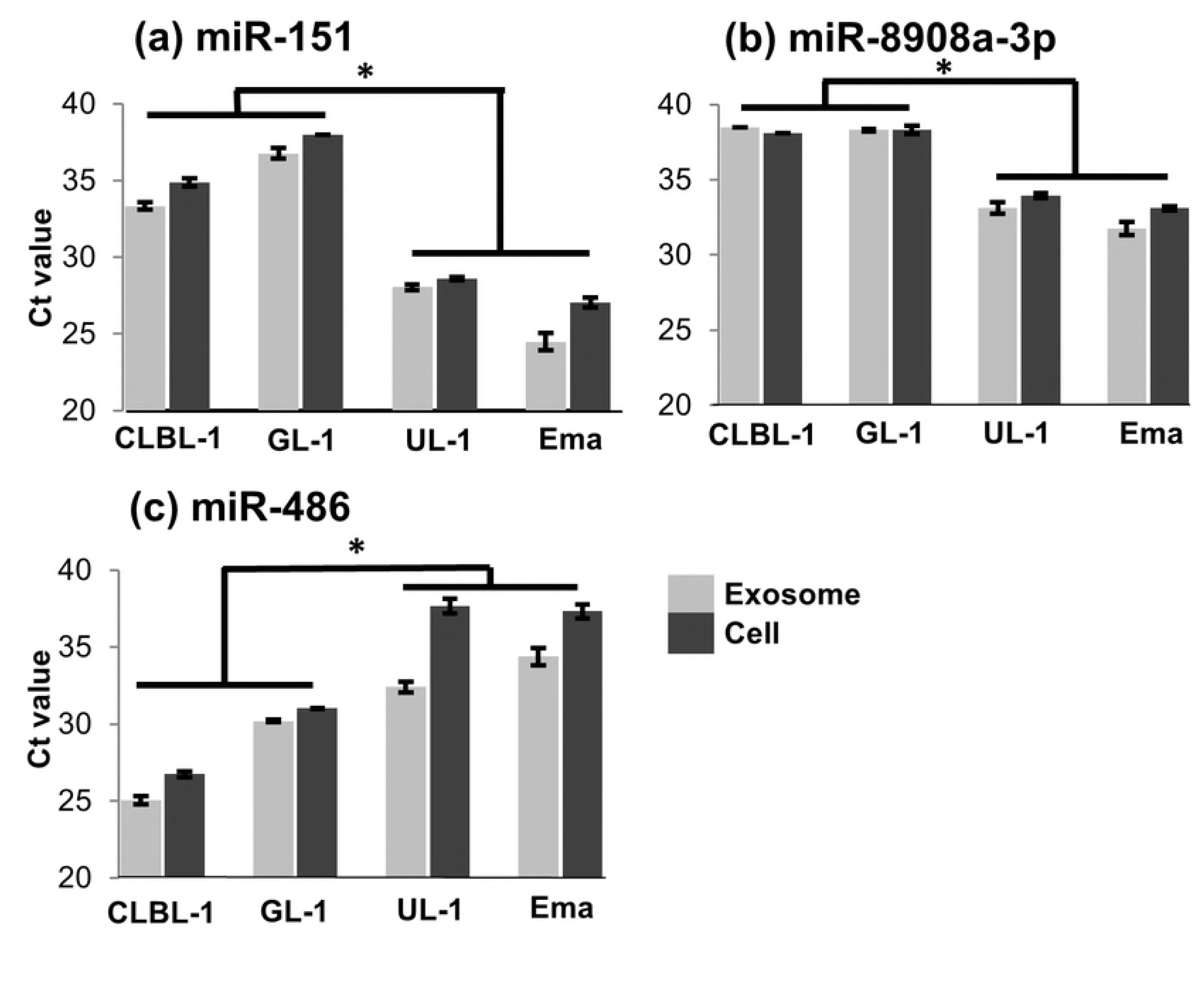
The separation of exosomal proteins from each cell line by SDS-PAGE. Lane M is the protein ladder. Lanes 1-4 correspond to the exosomal protein from CLBL-1, GL-1, UL-1, and Ema, respectively, and lanes 1’-4’ correspond to exosomal protein precipitated with trichloroacetic acid from CLBL-1, GL-1, UL-1, and Ema, respectively.

**S6 Fig.**
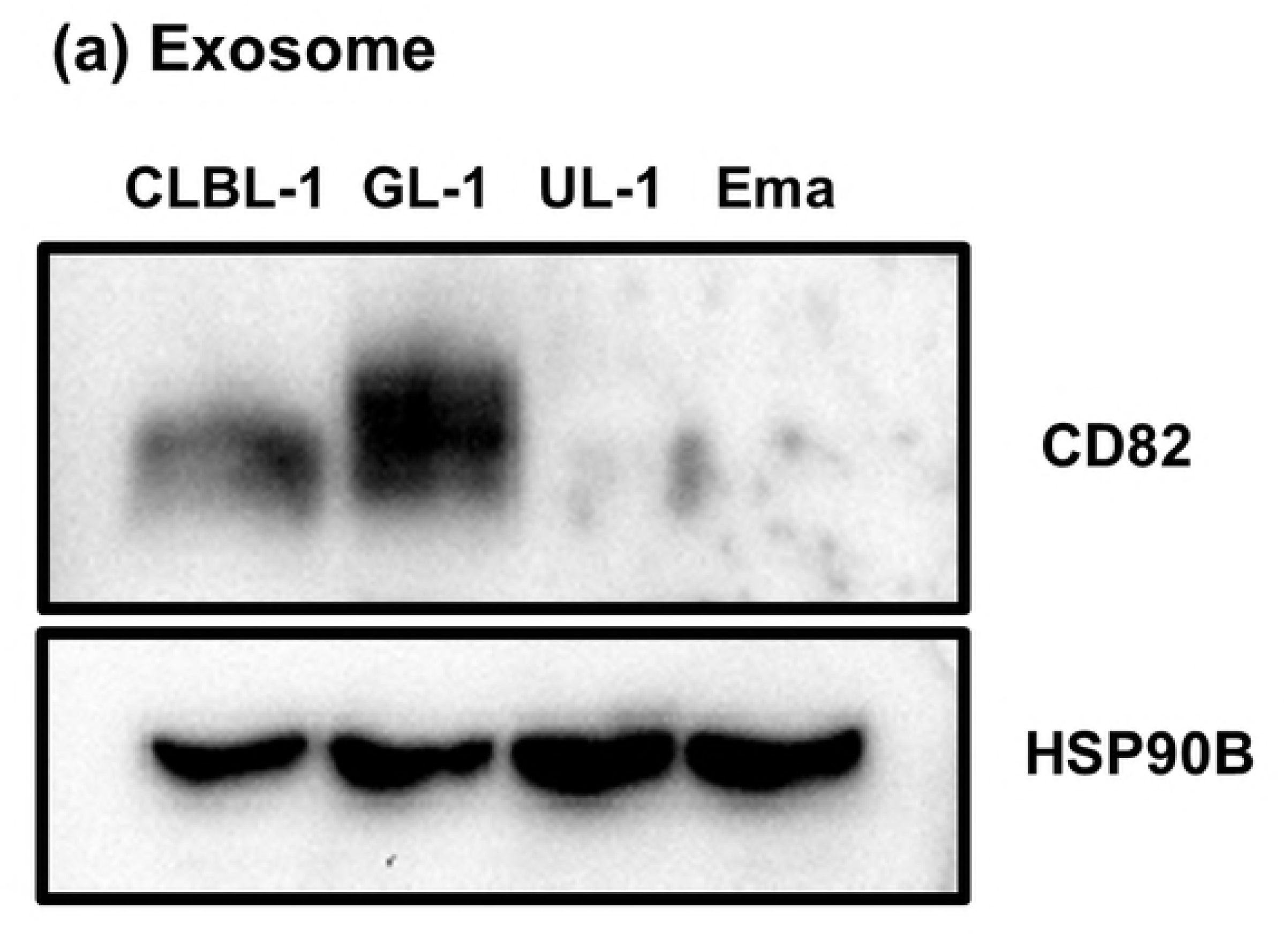
Heat maps showing the miRNAs whose amounts were significantly different between VCR-S cell lines and the VCR-R cell line in exosomes (a) and cells (b). In exosomes, the amounts of 11 miRNAs were significantly lower in VCR-S cell lines than in the VCR-R cell line, and those of 5 miRNAs were significantly higher in VCR-S cell lines than in the VCR-R cell line. In parent cells, the amounts of 8 miRNAs were significantly lower in VCR-S cell lines than in the VCR-R cell line, and those of 7 miRNAs were higher in VCR-S cell lines than in the VCR-R cell line.

**S7 Fig.**
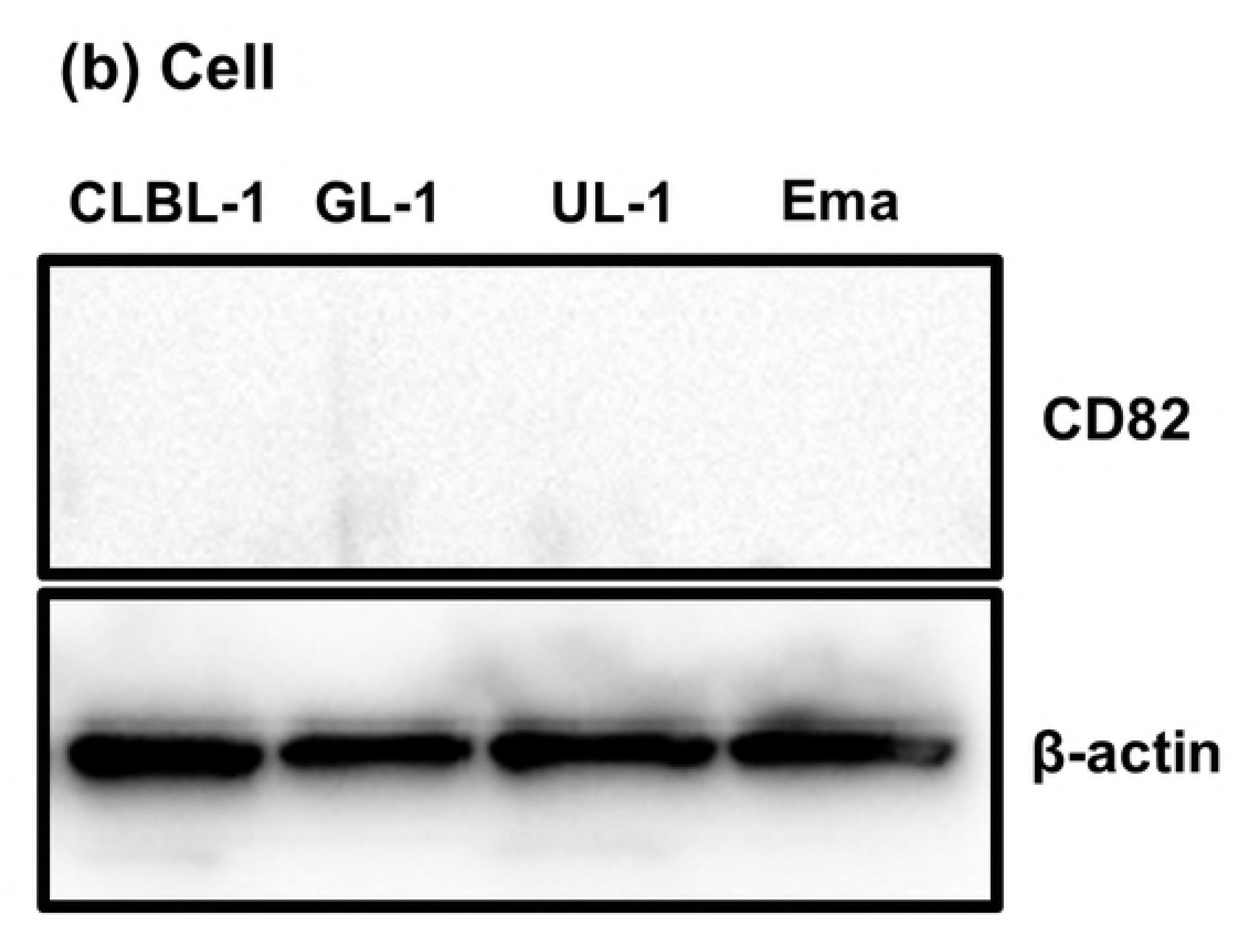
Full-length figures of blotting membrane that were used for the detection of CD82 (a, b), HSP90B (c), and β-actin (d) by Western blotting. The figures of the same membrane were shown in (a) and (b), but exposure time was different between these figures. In Fig 5, the figures of detection of CD82 within exosomes and parent cells were cropped from the different parts of (b). The figures of detection of HSP90B and β-actin were cropped from (c) and (d), respectively.

